# Massively parallel reporter assays for *CYP3A4* enhancer variants alongside their native promoter

**DOI:** 10.64898/2026.04.22.719677

**Authors:** Yelena Guttman, Beniamin Krupkin, Nadav Ahituv

## Abstract

Massively parallel reporter assays (MPRAs) enable high-throughput functional testing of thousands of *cis*-regulatory elements (CREs). However, conventional MPRAs typically use short CRE fragments combined with minimal promoters, which limits their ability to capture native regulatory interactions. Here, we present a modified MPRA framework that assays CREs together with their target promoter. We apply this approach to *CYP3A4*, a major pharmacogene with extensive interindividual expression variability. We assayed 1,214 variants across six *CYP3A4* regulatory regions, integrating variants from global populations, cancer genomes, and archaic humans. Most variants exerted modest to no effects on regulatory activity, consistent with strong functional constraint at the *CYP3A4* locus. However, we identified a subset of variants with significant regulatory impact, including variants that could be associated with altered drug response and cancer severity. Together, these results provide a comprehensive functional map of *CYP3A4* regulatory variation and establish a generalizable MPRA strategy for studying enhancer-promoter interactions.

## Introduction

Variation in gene expression is a key driver of phenotypic diversity, disease susceptibility, and variable response to therapy among individuals. A substantial proportion of this variation can be attributed to genetic polymorphism in *cis*-regulatory elements (CREs), such as promoters, enhancers, and silencers^1–4^. These regulatory variants can alter transcription factor (TF) binding, chromatin accessibility, and enhancer-promoter interactions, ultimately resulting in profound differences in gene expression across individuals and tissues.

Cytochrome P450 family 3 subfamily A member 4 (*CYP3A4*) encodes an enzyme that metabolizes over 50% of all clinically prescribed drugs and constitutes approximately 80% of the total CYP content in the human intestine and over 30% in the liver^5,6^. *CYP3A4* expression varies over 100-fold among individuals^7^, contributing to large inter-individual differences in drug metabolism, efficacy, and toxicity. While biological and environmental factors, including age, sex, diet, and co-administered medications, undoubtedly influence *CYP3A4* expression, genetic variation is also thought to play a significant role. However, the extent to which *CYP3A4* expression variability is genetically encoded remains unclear. *CYP3A4* also represents an example of evolutionary adaptation to diverse environmental and dietary pressures, as it is involved in the metabolism of plant-derived toxins and dietary compounds^8^. Comparative genomics reveals that *CYP3A4* diversity is the product of gene duplication events^9^ and positive selection following the split of the chimpanzee and the human lineages^10,11^.

As CYP3A4 metabolizes endogenous compounds, such as steroid hormones and vitaminlJD, variations in its expression or activity can disrupt hormonal and vitamin homeostasis, potentially increasing cancer susceptibility^12,13^. For example, the promoter variant *CYP3A4*1G* (rs2242480) has been linked to increased breast cancer risk in individuals under 50, particularly in early-stage and HER2-positive cases^14^. These findings suggest that *CYP3A4* regulatory variants may influence cancer risk by altering the metabolism of sex hormones and other endogenous molecules. Additionally, many anticancer agents are metabolized by or require activation through CYP3A4, underscoring the broader clinical relevance of *CYP3A4* genetic variation in cancer susceptibility, progression and treatment^13^.

Six CREs regulate *CYP3A4* (**Fig. 1a**): 1) the *CYP3A4* promoter; 2) an CCAAT/Enhancer-Binding Proteins (C/EBP) responsive element (C/EBPRE) that is 5.7 kilobases (kb) upstream of the transcription start site (TSS)^15^; 3) the xenobiotic-responsive enhancer module that mediates inducible transcriptional regulation in response to xenobiotics (XREM; -7.2 kb from TSS)^16,17^; 4) the constitutive liver enhancer module (CLEM4; −10.5 kb from TSS)^18^; 5) the distal regulatory region (DRR; +91 kb from TSS)^19^; 6) the intestinal hepatocyte nuclear factor 4 alpha (HNF4A) responsive region (IHRR; -8.8 kb from TSS)^20^. Despite this detailed annotation, the functional consequences of regulatory variation within these elements remain largely unknown and were carried out primarily on an individual variant basis.

**Figure 1:**
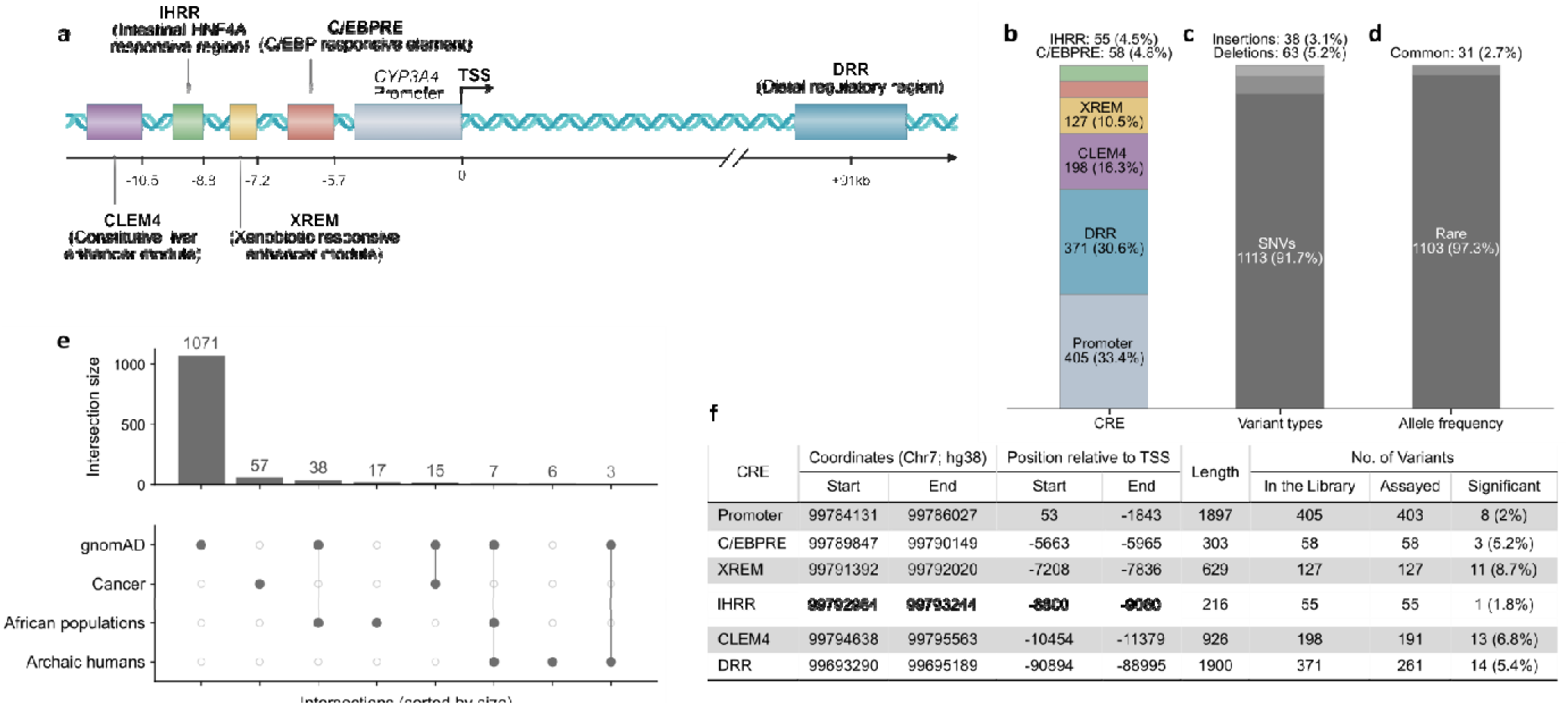
*CYP3A4 cis-*regulatory elements, variant library composition, and population-level representation across datasets. (**a**) Schematic representation of *cis-*regulatory elements (CREs) in the *CYP3A4* locus; (**b**) Distribution of variants across the five *CYP3A4* CREs; (**c**) Distribution of variants by types; (**d**) Allele frequency classification of variants based on gnomAD^28^ data (see Methods); (**e**) UpSet representation of intersections among gnomAD^28^, Cancer (TCGA)^29^, African populations, and Archaic datasets; (**f**) Genomic coordinates and variant summary for *CYP3A4* CREs.

Massively parallel reporter assays (MPRAs) can be used to functionally interrogate the impact of thousands of variants on CRE activity within a single experiment. By cloning candidate CREs alongside a transcribed barcode, MPRAs allow quantitative measurements of the transcriptional activity associated with each candidate CRE in a high-throughput manner^21^. MPRAs also provide a powerful framework for associating DNA sequence variation with transcriptional activity in a high-throughput and quantitative manner, and have been applied to dissect enhancer grammar and identify expression-modulating variants^22–24^. However, MPRAs have several limitations. One of the major limitations is that most MPRAs use short DNA sequences (typically 100-270 bp) as candidate CREs (cCREs). This is primarily due to the cost constraints of DNA synthesis, with oligonucleotide synthesis currently providing the most cost-efficient approach to obtain thousands of sequences. Previous studies have reported that the length of the tested sequence can affect MPRA results^24^. Moreover, the majority of MPRAs decouple enhancers from their endogenous target promoters, instead pairing them with a minimal promoter. While this approach simplifies assay design, it can fail to capture important features of endogenous regulation, including enhancer-promoter compatibility and the potential for interactions between variants in both regions. Although some enhancers can activate a broad set of promoters, 10-50% of tested enhancers exhibit promoter selectivity in mammals^25,26^. This specificity may be governed by combinations of TF binding motifs or other contextual sequence features. Thus, conventional MPRA approaches may overlook key dynamics relevant for transcriptional gene regulation.

To overcome these limitations, we developed a modified MPRA framework that retains both the full-length enhancer and its native promoter. This approach combines fragment-based cloning to reconstitute the full-length CREs, with long-read sequencing to accurately associate each variant with its corresponding barcode. This design provides a more precise representation of endogenous regulatory interactions, allowing for the systematic analysis of variant effects within their target promoter. We applied this modified MPRA to *CYP3A4*, one of the most clinically significant genes in pharmacogenomics^27^. By systematically introducing variants into *CYP3A4* CREs and testing their effects using our adapted MPRA design, we mapped the functional architecture of these regulatory regions and their functional core. We annotated binding sites for canonical and other TFs less associated with *CYP3A4* regulation. By integrating the measured regulatory effects of these variants with their population-specific allele frequencies, we were able to highlight five variants, both common and rare, as potential actionable pharmacogenomic targets. We also found cancer-associated variants that lead to a significant change in activity, primarily in melanoma and colorectal cancers. Combined, our work introduces a novel MPRA technology that assays longer CRE sequences and their variants alongside their target promoter and identifies variants associated with pharmacological response.

## Results

### *CYP3A4* library design

To comprehensively assay naturally occurring *CYP3A4* regulatory variants, we constructed a variant library by integrating four data sources: 1) the gnomAD database that contains whole genome sequencing (WGS) data from 76,215 individuals across seven populations^28^; 2) WGS data from African populations (see Methods); 3) WGS data of 2,577 patients across 21 cancer types from the Cancer Genome Atlas (TCGA)^29^; 4) the genomes of Neanderthal and Denisovan archaic humans^30,31^ (see Methods). The library covered the five previously published *CYP3A4* enhancers and its promoter (**Fig. 1a-b**). In total, the library contained 1,214 genetic variants comprising 1,113 single-nucleotide variants (93%) and 101 short indels (7%) (**Fig. 1c**). Amongst these variants, 31 were common (allele frequency >1% in at least one population) and 1,103 were rare (<1% in all populations) according to gnomAD (**Fig. 1d**), 57 cancer-specific, 17 African- specific, and 6 archaic-specific variants (**Fig. 1e**). Across all loci, gnomAD variant densities were distributed uniformly (Kolmogorov–Smirnov FDR>0.05) and Poisson-based enrichment analysis did not identify any hotspots or coldspots within individual loci.

### Optimization of baseline regulatory activity

To establish optimal assay conditions for the MPRA, we first assessed the baseline activity of reference hepatic (C/EBPRE, DRR, XREM, and CLEM4) and intestinal (IHRR) *CYP3A4* regulatory elements in human hepatocellular carcinoma (HepG2) and colorectal adenocarcinoma (LS174T) cells, respectively, using luciferase reporter assays. To identify the most suitable promoter configuration, each enhancer was tested upstream of three promoter constructs: 1) a minimal promoter commonly used in MPRAs; 2) the proximal 415 bp *CYP3A4* promoter; 3) the full-length 1.9 kb *CYP3A4* promoter. The proximal promoter produced the strongest transcriptional response, whereas the full-length promoter yielded lower activity, suggesting the presence of distal repressive element/s within the extended promoter region (**Supplementary Fig. 1a**). As saturation MPRA assays require robust regulatory activity to observe both gain and loss of function mutations, we chose the proximal promoter for our subsequent assays.

While HepG2 and LS174T are the most common models to assay the pharmacogenomic relevance of hepatic and intestinal *CYP3A4* expression, respectively, they require define the minimal transcriptional context necessary for *CYP3A4* CRE function, we co-transfected each enhancer with the TFs previously reported to regulate its activity, as well as with TFs known to modulate *CYP3A4* expression more broadly. Specifically, C/EBPRE was co-transfected with a plasmid expressing the CCAAT enhancer binding protein beta (*C/EBPB*; also called the Liver Activator Protein (*LAP*)), a potent *C/EBP* isoform that promotes *CYP3A4* transcription. The CLEM4 enhancer was assayed with the nuclear receptor subfamily 1 group I member 2 (*NR1I2*; also called the pregnane X receptor (*PXR*)); the DRR enhancer with *PXR* and hepatocyte nuclear factor 4 alpha (*HNF4A*); the XREM enhancer with *HNF4A*, the nuclear receptor subfamily 1 group I member 3 (*NR1I3*; also called the constitutive androstane receptor (*CAR*)), and *PXR*; and the IHRR enhancer in LS174T cells with *HNF4A*, *CAR,* and *PXR*. Because XREM is strongly activated via drug induction, XREM assays were performed both with and without rifampin, a potent antibiotic, provided 24 hours after transduction. Among the tested combinations, *LAP* co-expression with the C/EBPRE, PXR co-expression with the XREM enhancer and *HNF4A* co-expression with the IHRR led to increased transcriptional activity (**Supplementary Fig. 1b-f**). Interestingly, *PXR* co-expression reduced the transcriptional activity driven by both CLEM4, DRR and IHRR. Likewise, *HNF4A* and *CAR* modestly suppressed DRR- and XREM-mediated activation, respectively, suggesting that nuclear receptors may interfere with each other’s regulatory complexes or compete for overlapping binding partners. Rifampin treatment induced an over 100-fold increase in XREM-driven expression (**Supplementary Fig. 1g**), confirming its responsiveness to *PXR*-mediated drug induction. These findings are consistent with previous reports characterizing these regulatory regions^15–20^.

Based on our aforementioned results, all subsequent MPRA experiments incorporated the necessary co-factors to ensure robust enhancer activation: *LAP* co-expression for C/EBPRE, *HNF4A* co-expression for IHRR in LS174T cells, and *PXR* co-expression for XREM, with rifampin treatment included for XREM assays to maximally activate the *PXR* pathway. To provide robust co-factor expression, we generated HepG2 and LS174T stable lines expressing *PXR* and *HNF4A*, respectively, using lentiviral transduction. Since stable overexpression of *LAP* inhibits cell proliferation^34^, the generation of *LAP*-expressing stable lines was not feasible. Instead, *LAP* was transiently co-transfected with the C/EBPRE MPRA library in relevant assays.

### MPRA with native promoter and full-length enhancer

We next designed our MPRA cloning strategy and reporter constructs with two main objectives: 1) to assay variants within the full-length CREs; and 2) to measure CRE activity within the context of the native *CYP3A4* promoter, ensuring that enhancer-promoter compatibility is preserved. To achieve the first goal, we applied a fragment cloning approach: first, wild-type CREs were cloned into a pUC19 plasmid, a small (∼2.7 kb) high-copy-number plasmid chosen for its compact size, which simplifies whole-plasmid PCR amplification and contains an ampicillin resistance gene for selection. Next, synthetic segments of 200–300 bp containing all the mutations in that segment were cloned into the reference backbone plasmid. Finally, all segmented sub-libraries were pooled together to get a full set of variants for each CRE (**Fig. 2a**). The libraries were then amplified from the pUC19 plasmid backbone using PCR and cloned into an MPRA expression plasmid, pGL4-GFP (**Fig. 2b**, **Supplementary Data 3**). This strategy was used for long CREs (DRR and promoter) or when low sequence complexity did not allow direct synthesis (XREM). C/EBPRE, CLEM4 and IHRR sub-libraries were short enough to be ordered as full-length synthetic fragments. To achieve the second goal, having the CRE target promoter, we modified the MPRA construct to include the native *CYP3A4* proximal promoter, which has the most robust activity with *CYP3A4* CREs. Fifteen basepair (bp) barcodes were introduced by PCR downstream of the promoter. The enhancer library and barcoded promoter were assembled into the MPRA plasmid using Gibson assembly. We also generated an MPRA variant library of the *CYP3A4* promoter alongside the CLEM4 constitutive enhancer, as the promoter by itself does not provide sufficient regulatory activity that would allow to assay variant effects. Scrambled sequences of the CLEM4 and C/EBPRE CREs were used as negative controls for the experiment. In total, we tested 1,214 sequences across six CREs. All libraries, except for IHRR, were assayed in HepG2 cells; the IHRR library was assayed in LS174T cells. To maximize the dynamic range, MPRA libraries were introduced episomally at high copy number using the pGL4 backbone, which provides an increased dynamic range compared to the lentiMPRA method^35^.

**Figure 2:**
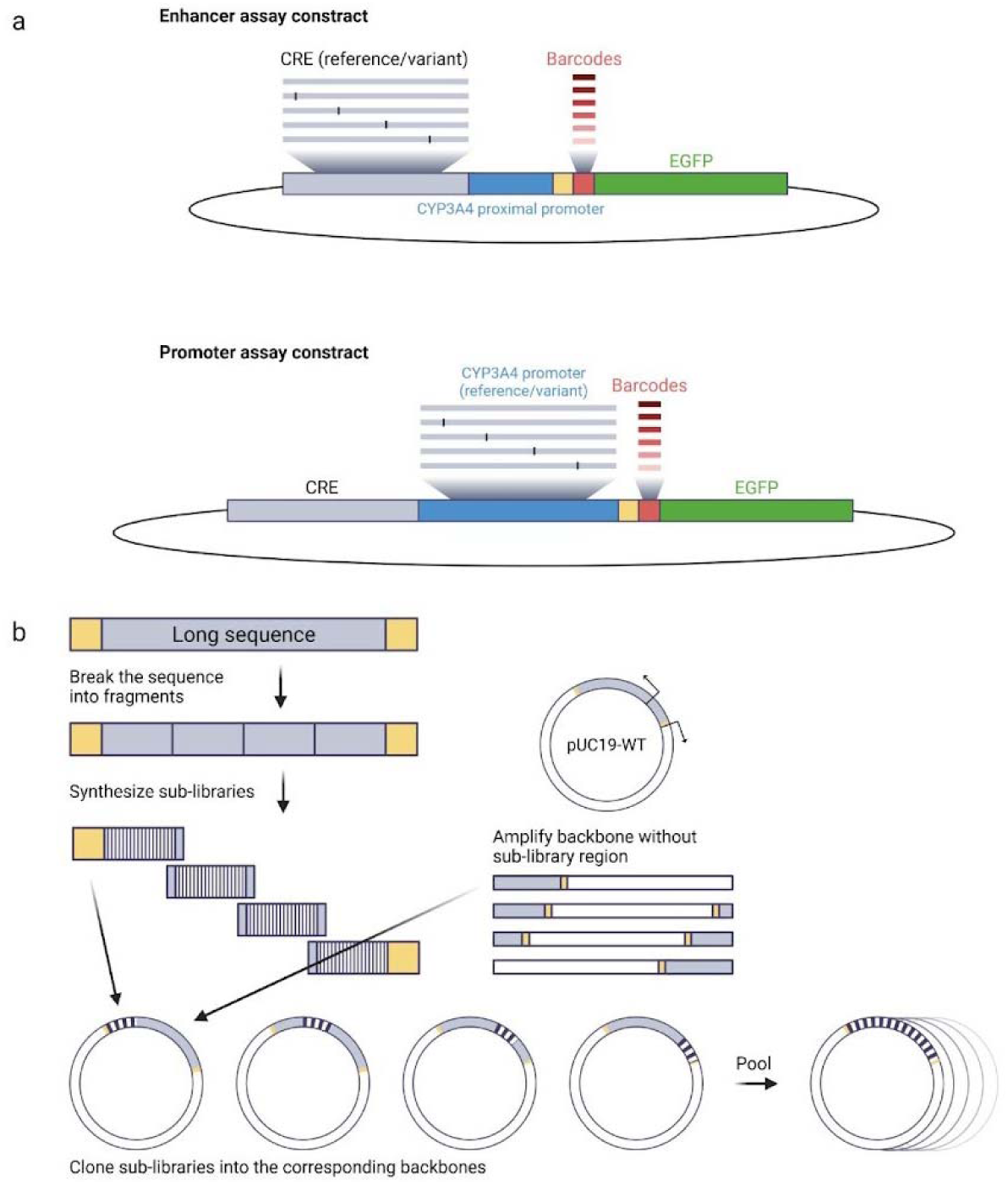
MPRA cloning strategy scheme. (**a**) Construct structures of CREs and promoter libraries. Enhancer reference/variant libraries are cloned upstream of the *CYP3A4* proximal promoter, barcodes and EGFP reporter gene. (**b**) Schematic representation of the fragment cloning strategy, where synthetic segments of 200–300 bp containing the desired variants in that segment are cloned into the reference backbone plasmid.

### MPRA identifies variants modifying *CYP3A4* CRE activity

Cells were transfected with the MPRA library using the aforementioned conditions for each CRE. DNA and RNA were harvested from cells, and barcodes were sequenced and analyzed using MPRAsnakeflow^36^ (see Methods). We had an average of 109 barcodes per variant and the correlation between replicates was >0.75 (**Supplementary Fig. 2**). Scrambled-sequence controls exhibited two orders of magnitude loss of activity, confirming that the assay accurately captures the intrinsic regulatory potential of the native CREs (**Supplementary Fig. 3**). Overall, amongst the 1,214 tested variants, 1,173 (97%) had robust MPRA activity scores. Of the 1,214 analyzed variants in the six *CYP3A4* CREs, 451 (38%) were identified as causing significant changes relative to the wild-type CRE (adjusted p-value<0.05), with the majority of significant effects being of small magnitude. Amongst them, 43 variants exceeded a 1.4-fold change threshold (log2 fold change (LFC)>0.5), of which 14 (33%) increased expression (by a median of 63%) and 29 (67%) decreased expression (by a median of 41%). The overall bias toward repressing variants, combined with our TF binding analysis, suggests that *CYP3A4* CRE single-nucleotide substitutions and short indels more often disrupt TF binding than create new functional activating sites. Significant variants were detected across all six CREs, with frequencies of 7.9% in XREM, 6.2% in CLEM4, 5.2% in C/EBPRE, 3.0% in DRR, 1.7% in the promoter fused to the CLEM4 constitutive enhancer and 1.8% in IHRR (**Fig. 1f**).

### Population-specific regulatory effects of *CYP3A4* variants

The vast majority of variants within *CYP3A4* regulatory elements are rare. However, because allele frequencies vary across genetic ancestry groups, we evaluated variants at the population-specific level to assess their potential pharmacogenomic relevance rather than applying a single global frequency cutoff. The most prominent candidate is the 99,795,181:C>G substitution in CLEM4, which is common in the African/African American population according to gnomAD (allele frequency 2.2%, with 14 homozygous individuals recorded) and is present at a lower frequency in the Latino/Admixed American population (0.39%). In our MPRA, this variant reduced regulatory activity by approximately 35%. Another notable variant is 99,694,580:T>C in the DRR, which significantly downregulated expression (LFC = -1.15). Although classified as rare, it is not negligible, with allele frequencies ranging from 0.2% to 0.4% across several populations, including European (non-Finnish), South Asian, and Latino/Admixed American groups. Similarly, 99,694,513:A>G, also located in the DRR, exhibited reduced activity (LFC = -0.57) and occurs at an allele frequency of 0.29% in the South Asian population. Three additional variants merit consideration due to their significant activity: 99,784,213:A>G in the promoter (LFC=-0.70), 99,791,812:T>C in XREM (LFC=1.33), and 99,795,146:C>T in CLEM4 (LFC=-0.58), with max allele frequencies of 0.021% in Europeans, 0.029% in Africans/African Americans and 0.013% in Europeans, respectively. The 99,694,424:C>T variant, although common across multiple populations and being the major allele in African/African American individuals, with allele frequencies up to 58% produced only a modest regulatory effect, below the cutoff of significance. Worth noticing that the 99,784,473:C>T, also known as the *1B allele, did not affect the expression in our MPRA (**Fig. 4a**, **Supplementary Data 1**).

### Cancer-associated variants

The majority (79%) of cancer-associated *CYP3A4* variants detected in the TCGA^29^ Databases were not observed in the general healthy population, as annotated by gnomAD^37^, suggesting that they represent rare or tumor-acquired mutations (**Fig. 1e**). Overall, cancer-associated variants exhibited modest but measurable regulatory effects (**Supplementary Data 2**). One substitution within the DRR exceeded the predefined significance threshold (|LFC| = 0.50), variant 99,693,619:T>G. Interestingly, this variant was found to be associated with hepatocellular carcinoma (HCC) in a whole-genome sequencing study of 321 tumors from 159 Japanese patients^38^. Other cancer-derived variants demonstrated lower than threshold but notable effects on expression, with several variants predominantly from melanoma and adenocarcinoma, but also one originating from brain cancer, exhibiting both up- and downregulatory impact (**Fig. 3c**). Notably, the repressive effects observed in melanoma- and colon-associated variants align with prior reports of reduced *CYP3A4* expression in these cancers^39–41^, supporting a potential contribution of noncoding regulatory variation to tumor-associated *CYP3A4* dysregulation.

**Figure 3:**
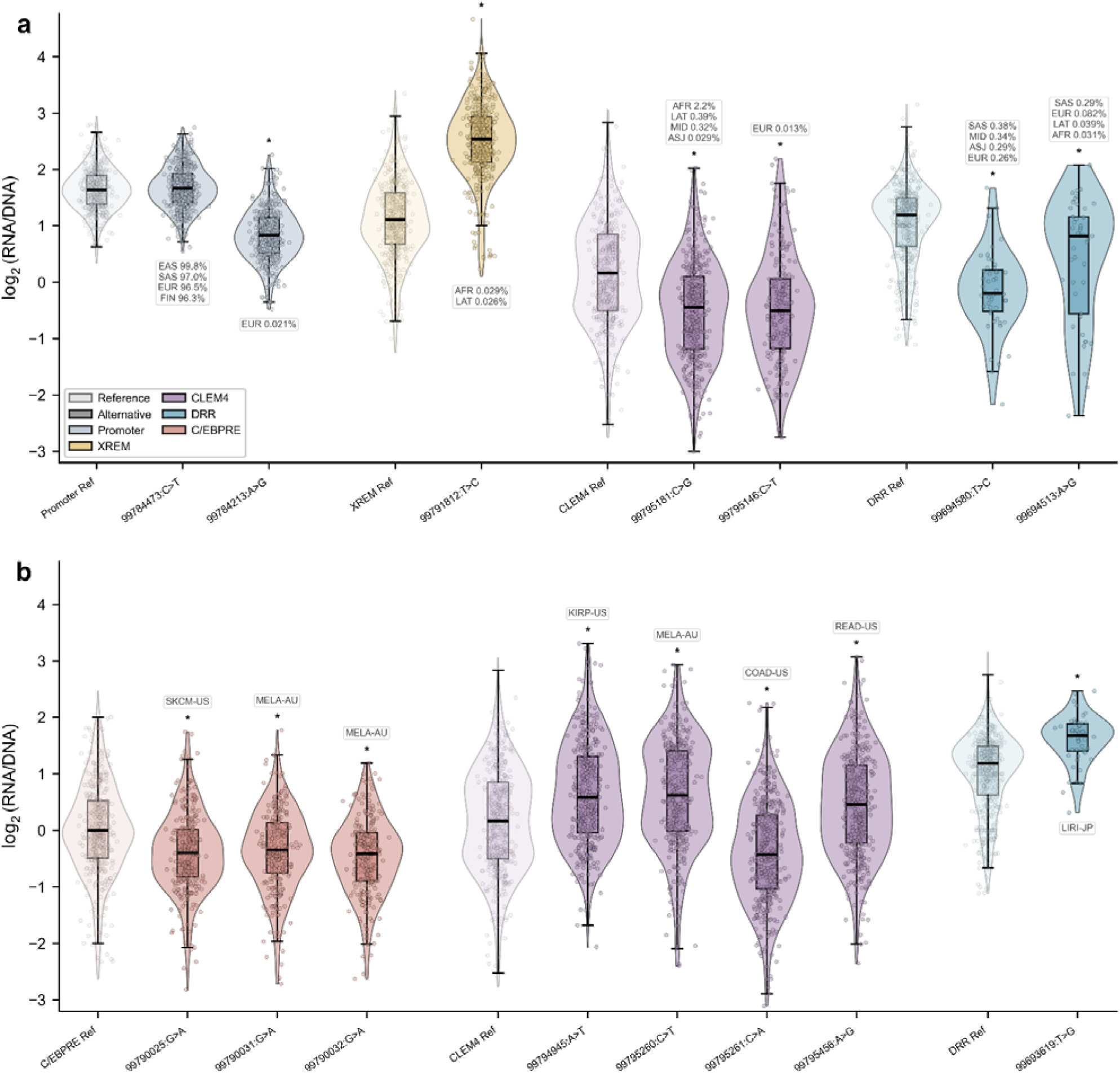
*CYP3A4* variant barcode activity distributions. Violin plots show per-barcode log₂(RNA/DNA) activity for reference and alternative alleles; asterisks indicate BCalm FDR < 0.05. (**a**) Top pharmacogenomic candidate variants with allele frequency in at least one population>0.01%. AFR, African/African American; ASJ, Ashkenazi Jewish; EAS, East Asian; EUR, European (non-Finnish); FIN, European (Finnish); LAT, Latino/Admixed American; MID, Middle Eastern; SAS, South Asian. (**b**) Cancer-associated somatic variants. COAD-US, colon adenocarcinoma (US); KIRP-US, kidney renal papillary cell carcinoma (US); LIRI-JP, liver cancer (Japan); MELA-AU, melanoma (Australia); READ-US, rectum adenocarcinoma (US); SKCM-US, skin cutaneous melanoma (US).

### *CYP3A4* promoter variant effects

The promoter significant variants were largely concentrated near the transcription start site (TSS) (**Fig. 4a**). Notably, the two activating promoter variants, located at positions 99,784,256 (all coordinates are from hg38) and 99,784,260 (LFC = 0.84 and 0.57, respectively), map to a previously reported silencer element^42^, containing two E-box motifs and a CCAAT box. Consistent with the removal of repression, TF binding site (TFBS) analysis indicated that the 99,784,260:A>G variant disrupts the E-box Snail family transcriptional repressor (SNAI) binding site, resulting in increased promoter activity. The 99,784,256:G>C variant leads to increased expression and gains a zinc finger 66 (ZNF66) TFBS. Several variants located closer to the TSS exhibited strong repressive effects (LFC = -0.62 to -1.3). Variants at positions 99,784,250 and 99,784,246 are located within the odd-skipped related transcription factor 1 (OSR1) binding site, disrupting the OSR1 motif and introducing a weak RE1-silencing transcription factor (REST)^43^ motif which represses neuronal genes in non-neuronal tissues, respectively. A closely located variant containing 25 (ZSCAN25) TFBS. Although *ZSCAN25* overlaps the *CYP3A* locus on chromosome 7, no evidence of regulatory involvement has been demonstrated so far^44^. Additional downregulating variants at positions 99,784,209 and 99,784,213 disrupted TFBS recognized by members of the Forkhead box (FOX) family hepatic TFs^45^ (**Fig. 4b**).

**Figure 4:**
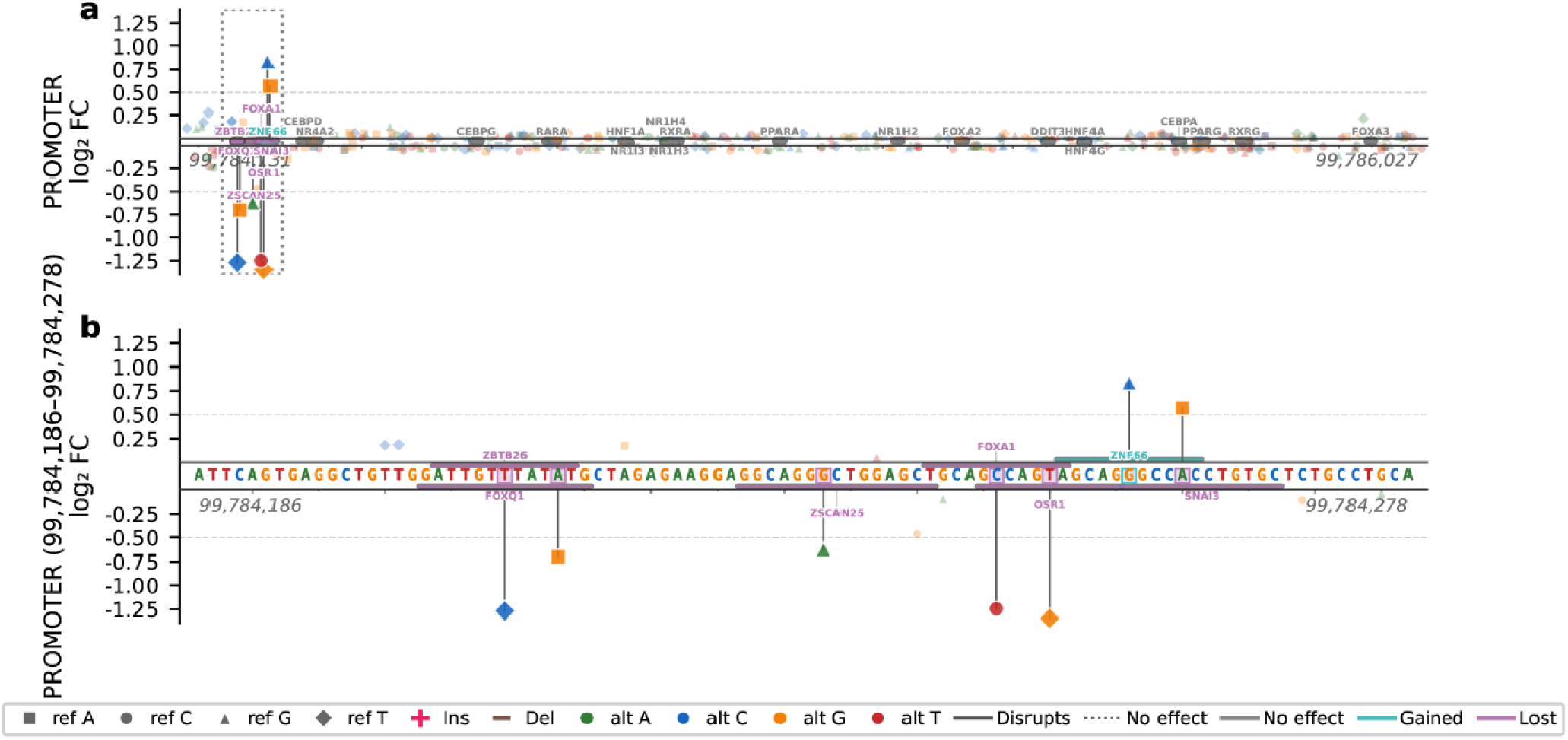
Effects of variants in the *CYP3A4* promoter. (**a**) Log_2_ fold change for all tested variants, ordered by genomic position within the *CYP3A4* promoter. Predicted transcription factor binding motifs from JASPAR are shown as bars aligned to the sequence. Motifs present in the reference sequence are shown in gray, variant-induced motif gains in teal, and motif disruptions caused by variants in purple. The functionally variable region is indicated by a dashed box. (**b**) Expanded view of the functionally variable region highlighted in (a), displayed using the same format. Genomic coordinates are provided for hg38.

### *CYP3A4* CRE variant effects

Only three variants significantly reduced C/EBPRE activity. The strongest effect was observed for the 99,789,956:A>G variant (LFC = -0.76), which disrupted predicted TFBS for activating transcription factor 4 (ATF4) and for the C/EBP family, the TFs that gave this CRE its name. ATF4 partners with C/EBPs to form active heterodimers^46^. No substantial transcriptional effect was found for variants located within other canonical liver TFBS such as C/EBPA, C/EBPG, NR1I3, and Peroxisome proliferator-activated receptor (PPAR) family (**Fig. 5a**)

**Figure 5:**
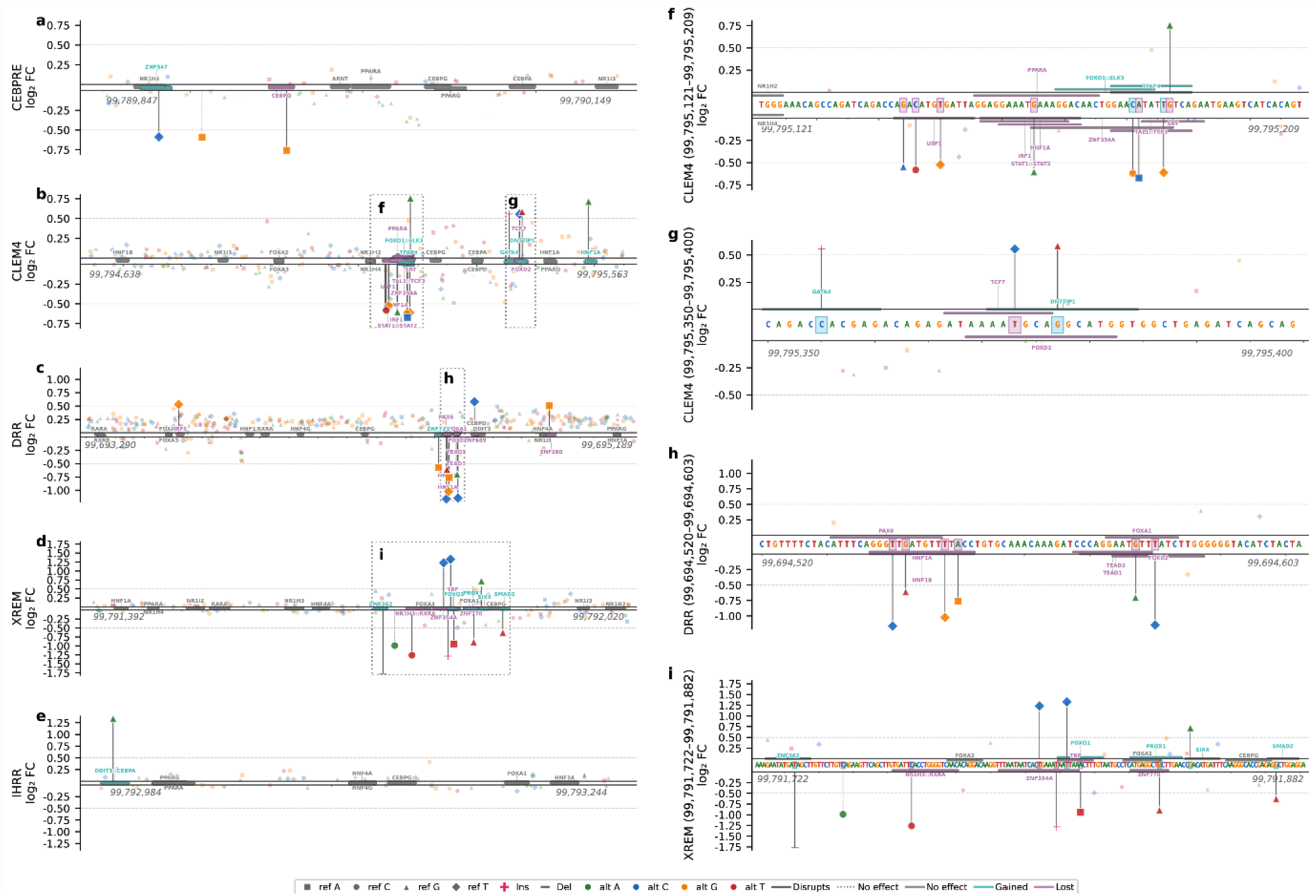
Effects of variants across *CYP3A4* CREs. (**a-e**) Log_2_ fold change for all tested variants, ordered by genomic position within the (**a**) CEBPRE, (**b**) CLEM4, (**c**) DRR, (**d**) XREM, and (**e**) IHRR. (**f-i**) Expanded views of functionally variable regions indicated by dashed boxes in panels (b–d): (**f**) CLEM4 region 1, (**g**) CLEM4 region 2, (**h**) DRR, and (**i**) XREM. Genomic coordinates are provided for hg38. Predicted transcription factor binding motifs from JASPAR are shown as bars aligned to the sequence. Motifs present in the reference sequence are shown in gray, variant-induced motif gains in teal, and motif disruptions caused by variants in purple.

Within the CLEM4 element, we identified twelve variants with significant regulatory effects, seven that reduced enhancer activity (LFC = -0.52 to -0.67) clustered within a narrow region spanning 99,795,144-99,795,186. Five variants that increased enhancer activity are located downstream (LFC = 0.55 to 0.76), three of them localized to 99,795,355-99,795,377 (**Fig. 5b,f,g**). The activating module harbors a USF1 TFBS. USF1 loss, caused by three different variants, resulted in downregulation, consistent with a previous report on the CLEM4 enhancer^18^. The variant at position 99,795,165 overlapped HNF1A and lowered the motif matching score. Alongside the loss of activating TF binding sites, downregulation can also be linked to the gain of binding repressors such as the FOXO1::ELK3 pair. In addition, four variants within a 7-bp region (99,795,181-99,795,188) overlapping two bHLH motifs variants increasing affinity towards TAL1::TCF3 TFBS had a decrease of expression, while the variant that decreased affinity increased expres ion, suggesting a repressive role at this locus. The introduction of additional HNF1A and GATA TFBS by variants 99,795,490:G>A and 99,795,355:C>CACGAGACAGAGATA, respectively, was associated with increased activity.

Within the DRR element, the majority of significant variants reduced regulatory activity relative to the reference sequence. Most of these loss-of-function substitutions clustered between 99,694,540-99,694,580 (LFC = -0.52 to -1.40), suggesting that this interval represents a functionally variable core of the DRR with both HNF1A/B and FOX family TFBS. As expected the enhancers. In addition, a TEAD disrupting variant reduced expression. While not part of the canonical regulation, the YAP/TEAD signalling pathway was recently shown to regulate *CYP* induction^47^ (**Fig. 5c,h**).

Within the XREM region, ten variants exhibited a significant change in regulatory activity in the interval spanning 99,791,733-99,791,873, suggesting the presence of a functionally variable subregion within XREM. Most variants reduced enhancer activity, with the strongest effect observed for the 23-bp deletion variant at position 99,791,733, which decreased expression by approximately 1.8-fold. The deleted region contained low-complexity ZNF TFBS and weak FOX family affinity sites. In contrast, three variants (99,791,804, 99,791,812, and 99,791,848) increased regulatory activity, resulting in approximately 1.2-, 1.3- and 0.7-fold upregulation, respectively, possibly due to the gain of binding sites for FOX family TFBS. The variant 99,791,839:G>T introduced a predicted Prospero-Related Homeobox 1 (PROX1) binding site; PROX1 has been shown to interact with PXR^48^, forming a complex that inhibits PXR-mediated activation of *CYP3A4*. Other variants led to a reduction in enhancer activity but without explanatory TFBS perturbations. Notably, none of the significant variants disrupted predicted TFBS for PXR, suggesting selection preserving canonical regulation (**Fig. 5d, i**).

A single variant, 99,792,990:G>A, significantly increased IHRR activity (LFC = 1.33), gaining a weak affinity site for DNA damage inducible transcript 3 (DDIT3, also known as C/EBP homologous protein (CHOP))::C/EBPA binding, both are C/EBP family TFs. Despite the observed decrease in IHRR activity upon PXR co-expression in luciferase assays, no predicted PXR binding sites were identified within this region, suggesting that the inhibitory effect may arise from competition for shared binding partners rather than from direct PXR binding (**Fig. 5e**).

### Archaic human variant effects

For the six variants we selected using a comparative analysis of archaic and modern human genomes, none exceeded the predefined MPRA activity threshold, with all variants showing absolute log fold changes below 0.25. We additionally identified ten variants shared between archaic and modern genomes, indicating that these alleles predate the modern-archaic split. Similar to the archaic-specific variants, these shared variants also did not exceed the MPRA activity threshold (|LFC| < 0.3). In present-day human populations, the shared variants are individually rare and are not observed together as a common haplotype, whereas in archaic data, derived from single reference genomes, they co-occur within individuals (nine variants in Neanderthal and seven variants in Denisovan (**Fig. 1e**). These variants may represent rare remnants of ancestral regulatory states, while modern populations predominantly carry derived configurations that better fit contemporary selective constraints.

### Functional analysis of DRR haplotypes

As our MPRA tested only single variants, we next wanted to test the effect of haplotypes in these CREs on regulatory activity. We determined haplotype inference based on shared allele frequencies across global populations and linkage disequilibrium data from the LDHap database^49^. This revealed two multi-variant haplotypes located within the DRR enhancer. These included the double haplotype: 99,694,578:T>G (rs544295692) and 99,694,579:T>C (rs562971068), and the four variants haplotype: 99,694,424:C>T (rs776744), 99,694,585:GG>G (rs11353593), 99,694,756:CACA>CA (rs61017966), and 99,695,131:C>T (rs776742). The double haplotype is rare, observed at maximal frequencies of 0.34%, 0.29%, and 0.26% in Middle Eastern, Ashkenazi Jewish, and European populations, respectively. The four variant haplotype, on the other hand, is relatively common, with allele frequencies of 22% in Latino, 28% in East Asian, 34% in South Asian, and 21% in Middle Eastern populations, compared to 9% in Europeans and a major allele frequency of 58% in Africans.

To assess potential epistatic interactions between co-occurring variants, we conducted luciferase reporter assays comparing the regulatory activity of each haplotype to that of its constituent single variant. This allowed us to assay the regulatory activity of two variants for which MPRA scores were not determined: 99,694,579:T>G and 99,694,585:TG>T. Consistent with the MPRA results, 99,694,580:T>C significantly decreased activity, whereas variants 99,694,424:C>T, 99,694,756:CCA>C, and 99,695,132:C>T of the four-variant haplotype deviated from the MPRA results and showed moderately reduced activity (**Supplementary Fig. 4**). In both haplotypes, the combined regulatory effect of co-occurring variants was less than expected for a purely additive interaction, suggesting antagonistic interactions among variants. In the double haplotype, the variants occur in consecutive nucleotides, which may contribute to a redundant effect, whereas in the four-variant haplotype, the variants are spread across 707 bp, suggesting that the binding of different factors might be affected.

## Discussion

In this study, we developed and applied a modified MPRA framework that preserves both full-length CREs and their native promoter to systematically interrogate regulatory variation at the *CYP3A4* locus. By assaying more than 1,200 naturally occurring variants spanning six regulatory regions, we provide a comprehensive functional map of noncoding variation affecting one of the most clinically important pharmacogenes. Our results refine long-standing assumptions about the genetic basis of *CYP3A4* expression variability, clarify the role of specific TFs that regulate it, and provide a modified MPRA framework that assays full-length CREs with their endogenous target promoter, which can be implemented to study variation across the genome.

Previous studies have demonstrated that enhancer-promoter compatibility is a key determinant of regulatory output^26^ and that classical episomal and lentiMPRA designs might not be as robust in detecting regulatory effects of weak or context-dependent variants, even in disease-associated CREs, with results influenced by promoter choice and assay configuration^22,35^. To mitigate these limitations, we matched the enhancers to their native *CYP3A4* promoter and optimized the transcriptional and environmental context, including the required TFs and chemical inducers, prior to the MPRA assay to ensure measurable baseline activity across all tested CREs. In our study, the majority of variants produced no detectable or only modest effects on regulatory activity, a pattern consistent with expectations under purifying selection and with the known robustness of essential detoxification gene regulation. Importantly, despite these constraints, we were able to detect regulatory effects exceeding a |LCF|=0.5 threshold in approximately 5% of the tested variants. This distribution is consistent with findings from saturation mutagenesis MPRAs of diverse enhancers^22,35^, in which most single-nucleotide substitutions exert subtle quantitative modulation rather than large disruptions of regulatory function.

Given the central role of CYP3A4 in the metabolism of endogenous compounds, dietary toxins, and pharmaceuticals, the *CYP3A4* locus has been shaped by positive selection that enabled adaptation to changing environments while simultaneously enforcing strong purifying constraints against highly disruptive mutations. Variants persisting in generally healthy populations, such as those represented in gnomAD, likely reflect survival bias toward functionally tolerated regulatory effects. Accordingly, the absence of measurable effects for many variants in canonical TF binding sites (e.g., LAP in C/EBPRE, HNF4A in IHRR, and PXR in XREM) likely reflects strong functional constraint and selective depletion of disruptive mutations, rather than a lack of regulatory importance.

Our variant-level analysis revealed regulatory logic extending beyond the canonical nuclear receptors classically associated with *CYP3A4*. Beyond PXR, CAR, and HNF4A, C/EBP and FOX family, which are essential mediators of inducible and basal regulation, functional variants in our MPRAs altered predicted binding sites of other TF families, including USF1, TEAD1 and TAL1. Recurrent disruption of USF1 motifs within the CLEM4 enhancer resulted in consistent downregulation, reinforcing its role as an activating factor and demonstrating that naturally occurring human variants can measurably modulate this activity. Similarly, variants affecting FOX family motifs repeatedly reduced enhancer activity across multiple CREs, underscoring the importance of Forkhead factors in maintaining hepatic regulatory output. Although *CYP3A4* expression is highest in the liver and intestine, low-level expression is observed in multiple extrahepatic tissues^49^, including the blood-brain barrier and skin, and is dynamically regulated during development, most notably through the postnatal switch from *CYP3A7* to *CYP3A4/5* expression. The enrichment of broadly acting bHLH and homeobox motifs within *CYP3A4* CREs suggests that these enhancers encode general regulatory grammar, permitting tissue- and stage-specific tuning of expression through differential TF availability rather than strict tissue-restricted control. It is worth noting that TF binding predictions were inferred from motif analyses rather than direct occupancy measurements, and thus, causal links between specific TFs and regulatory effects will require orthogonal validation using chromatin-based assays such as ChIP-seq or TF perturbations.

The *CYP3A4*1B* variant (rs2740574; 99,784,473:C>T) was extensively studied in GWAS and drug response studies. However, data associated with this variant increased the risk for cancer^50^ and increased pharmacokinetics of tacrolimus^51^ and opioids^52^ was inconclusive. It was therefore suggested that the association may reflect linkage disequilibrium with the functional variant *CYP3A5*1* rather than a direct effect of *CYP3A4*1B* on gene expression^53^. In support of this hypothesis, the 99,784,473:C>T variant did not exhibit a significant regulatory effect in our MPRA, consistent with a prior study that also demonstrated the lack of a direct transcriptional impact^54^. Our dataset also included eight previously reported *CYP3A4* regulatory variants cataloged in PharmVar^55^, none of which had prior direct functional validation. Functional assessment of these variants revealed minimal effects on regulatory activity, suggesting limited regulatory impact. Together, these findings support the long-standing missing heritability problem in *CYP3A4* regulation^56^. While genetic contributions have been widely hypothesized to underlie interindividual differences in enzyme levels, prior studies^57,58^ have noted that known variants explain only a small fraction of the up to 100-fold variability observed within individuals. Our data refine this notion by showing that common regulatory variants exhibit little to no measurable regulatory activity, whereas detectable effects are largely restricted to rare variants with modest effect sizes. This architecture is consistent with a regulatory system in which baseline genetic variation is constrained, and dynamic environmental and physiological inputs, including xenobiotic exposure, inflammation, and hormonal regulation, drive the majority of expression variability^59^.

Our MPRA library included cancer-associated variants located within *CYP3A4* CREs. One variant significantly increased CRE activity, while ten others showed noticeable effects that did not reach statistical significance. Several cancer-associated regulatory variants identified here originated from epithelial malignancies, including melanoma, colorectal adenocarcinoma, and hepatocellular carcinoma, where *CYP3A4* dysregulation has been repeatedly observed^39,60^. Given the role of *CYP3A4* in xenobiotic metabolism, detoxification, and activation, including anticancer drugs^40^, altered regulatory activity in these tumors may influence both endogenous metabolic states and response to anticancer therapies. The predominantly repressive effects observed for melanoma-and colorectal cancer-derived variants are consistent with prior reports of reduced *CYP3A4* expression in these malignancies^39,41^.

The long-term objective of this work is to identify previously overlooked pharmacogenomic candidates within *CYP3A4*, a locus for which clear genotype-guided prescribing recommendations are currently lacking despite its central role in drug metabolism. Among the variants evaluated, we identified a single regulatory substitution that produced a substantial reduction in activity in our MPRA and is also common within the African/African American genetic ancestry group. The combination of measurable functional effect and population-specific prevalence makes this variant a compelling candidate for further investigation aimed at establishing its potential impact on *CYP3A4*-mediated drug metabolism. In addition, several variants with strong regulatory effects were observed at lower allele frequencies. Although individually rare, variants with allele frequencies as low as ∼0.1% may still affect large absolute numbers of individuals when considered at the scale of global populations. Ultimately, clinical outcome data will be required to determine whether the variants characterized here ultimately influence CYP3A4-mediated drug metabolism and can be actionable pharmacogenomic targets.

Although our MPRA framework preserves full-length CREs together with their native promoter, the assays were performed using episomal reporter constructs and therefore do not fully recapitulate endogenous chromatin context, higher-order genome organization, or long-range regulatory interactions that may modulate *CYP3A4* expression *in vivo*. In some cases, we did not directly observe motif affinity to be explanatory for the expression difference. In these cases, the binding *in vivo* could be affected by variant flanking effects and variant DNA-shape modifications^61^. In addition, while HepG2 and LS174T cells are widely used hepatic and intestinal models, respectively, they cannot capture the full spectrum of tissue-specific, developmental, or physiological states in which *CYP3A4* is regulated, nor the dynamic environmental exposures known to strongly influence its expression. Notably, these cell lines also lack expression of several key TFs required for optimal *CYP3A4* regulation, necessitating their exogenous supplementation in our assays. Despite these limitations, the high reproducibility across biological replicates and the consistency of effect sizes observed across regulatory elements support the robustness of our conclusions regarding the overall architecture and constraints of *CYP3A4* regulatory variation.

## Methods

All restriction enzymes were purchased from New England Biolabs. All primers were ordered from Integrated DNA Technologies (IDT). All primer sequences are provided in **Supplementary Data 2**. NEBNext Ultra II Q5 Master Mix (New England Biolabs, M0544) was used for all library PCR amplification reactions.

### Library design

Enhancer and promoter coordinates were extracted from previous publications (**Fig. 1f**). Variant data of the six CRE regions was downloaded from gnomAD (v3.1.2)^37^ as VCF files based on 76,156 WGS mapped to the GRCh38 reference sequence. The archaic human data was downloaded from Prüfer et al (2014)^31^ as VCF files mapped to hg19 and was converted using Liftover^62^ to hg38. African populations dataset was created by extracting variants from WGS of 180 African individuals from 12 indigenous African populations^63^. Additional variants were identified from WGS of tumor samples from 2,577 individuals across 21 tissues obtained from open access release data of The Cancer Genome Atlas program (TCGA)^29^.

### Cell culture

HepG2 and LS174T cells (ATCC, #HB-8065 and #CL-188, respectively) were grown in Eagle’s Minimum Essential Medium (EMEM) (Corning, 10-009-CV) supplemented with 10% FBS (Gibco, A38400-01) and 1% Penicillin-Streptomycin (10,000 U/mL; Gibco, 15140-163). For HepG2 cells, cell culture plates were coated with Collagen coating solution, using 1 ml per 10 cm^2^ culture area (Sigma-Aldrich, 125-50) for 30 minutes at 37°C. LentiX 293T cells (Takara, 632180) were grown in Dulbecco’s Modified Eagle Medium (DMEM) containing Glutamax and pyruvate (Gibco, 10569069) and supplemented with 10% heat-inactivated FBS (Gibco, A38400-01). All cells were grown at 37°C with 5% CO_2_ and were regularly tested for the absence of mycoplasma contamination using a Mycoplasma PCR Detection Kit (Applied Biological Materials Inc., G238).

### Luciferase reporter assays

Reference sequences of the six CREs were synthesized by Twist Bioscience (XREM, C/EBPRE, and IHRR) or by Integrated DNA Technologies (promoter, CLEM4, and DRR) with homology arms for Gibson assembly. XREM, C/EBPRE, CLEM4, IHRR, and DRR enhancer fragments were cloned upstream of either a minimal promoter, the proximal *CYP3A4* promoter (415 bp), or the full-length *CYP3A4* promoter into the *SacI* and *NcoI* sites of the pGL4.23c vector (Promega) using the NEBuilder HiFi DNA Assembly Cloning Kit (New England Biolabs, E2621). Each enhancer-promoter construct, along with the empty pGL4.23c vector (negative control) and pGL4.13 vector (positive control), was co-transfected with pGL4.74 (Renilla luciferase control; Promega) into HepG2 cells (5×10^4^ cells/well in 24-well plates) using X-tremeGENE HP DNA transfection reagent (Sigma-Aldrich, XTGHP-RO) at a reagent:DNA ratio of 2:1, following the manufacturer’s protocol. For XREM constructs, transfected cells were treated 24 hours post-transfection with 10 µM rifampin (Sigma-Aldrich, R3501) or 0.1% DMSO (Sigma-Aldrich, D2650) as a vehicle control. Each condition was tested using three biological replicates. Firefly and Renilla luciferase activities were quantified using the Dual-Luciferase Reporter Assay System (Promega, E1910) on a GloMax microplate reader (Promega). Enhancer activity was calculated as the ratio of Firefly to Renilla luciferase signal, normalized to the corresponding control. To assess TF dependencies, cells were co-transfected with expression plasmids for pregnane X receptor (*PXR*), constitutive androstane receptor (*CAR*), hepatocyte nuclear factor 4 alpha (*HNF4A*), and liver-enriched activator protein (*LAP*). *PXR* and *LAP* expression vectors were obtained from VectorBuilder containing the human PXR or LAP coding sequence under the control of the EF1A promoter in a pLV lentiviral backbone (see **Supplementary Data**), while *CAR* and *HNF4A* plasmids were ordered from AddGene (#141652 and #141663, respectively).

### Generation of stable cell lines expressing supplementary TFs

The stable HepG2 cell line expressing PXR and the LS174T cell line expressing HNF4A were generated using a lentiviral transduction system. The same TF expressing vectors as were used for the luciferase reporter assays were used here. For lentivirus production, 3×10^6^ Lenti-X 293T cells were seeded in 15 cm dishes and cultured to 70-80% confluence. Cells were transfected with 6.4 µg of the PXR lentiviral plasmid and 12.8 µg of Lenti-Pac HIV mix (GeneCopoeia, LT001) using EndoFectin Lenti reagent (GeneCopoeia, EF001), following the manufacturer’s protocol. After 8 hours, the medium was replaced and ViralBoost reagent (Alstem, VB100) was added. Virus-containing supernatant was collected 48 hours post-transfection and concentrated using the Lenti-X Concentrator (Takara, 631231) according to the manufacturer’s protocol. HepG2 and LS174T cells were transduced with the concentrated lentivirus and, after 48 hours, subjected to puromycin selection (2 µg/mL) for 14 days. Surviving colonies were expanded and screened for integration using primers for the lentiviral WRPE sequence (primers WRPE-F, WRPE-R) and expression by qPCR using *TUBB* as a control gene (primer sequences for HNF4a-qpcr-F, HNF4a-qpcr-R, PXR-qpcr-F, PXR-qpcr-R are in **Supplementary Data 2**).

### MPRA

#### Cloning of pGL4-GFP as a backbone for MPRA

The GFP coding sequence was amplified from the pLS-mP vector (Addgene, #81225) using primers designed to add 5′ and 3′ overlaps homologous to the pGL4c vector digested with *HindIII* and *XbaI* (Primers GFP-F, GFP-R). Gibson assembly was performed using the NEBuilder HiFi DNA Assembly Master Mix according to the manufacturer’s protocol. The assembled pGL4-GFP product was transformed into *E. coli DH5*α competent cells (New England Biolabs, C2987) and positive clones were validated by plasmid sequencing (Plasmidsaurus).

#### Fragment cloning

The XREM, DRR, and *CYP3A4* promoter libraries were assembled using a fragment-based cloning strategy. Reference CREs were PCR-amplified (primers pUC19-CRE-WT-F, pUC19-CRE-WT-R) to introduce overhangs compatible with the pUC19 vector digested with *HindIII* and *XbaI*. To enable efficient incorporation of variants across the full length of each CRE, the libraries were divided into three (XREM and Promoter) or four (DRR) sub-libraries, each covering a distinct subregion. Variant-containing sub-libraries were synthesized by IDT (Promoter, and DRR fragments 2-4), or TWIST (DRR fragment 1 and XREM) with homology arms compatible with the remaining plasmid sequence. For each sub-library, the complementary pUC19-reference CRE backbone was linearized (**Supplementary Data 2**) and PCR-amplified (primers pCU19-CRE-F#-F, pCU19-CRE-F#-R), and Gibson assembly was used to insert the variant sub-library into the plasmid. Assembly products were purified with 0.65X Ampure XP beads (Beckman Coulter, A63881), and 34 ng of assembly product was transformed into 50 μL *E. coli 10-beta* electrocompetent cells (New England Biolabs, C3020) using a Gemini X2 electroporator (settings [voltage: 2 kV; resistance: 200 Ohms; capacitance: 25 uF; number of pulses: 1; gap width: 1mm]). Colonies were grown overnight on ampicillin plates and subsequently processed using a Midiprep Kit (Qiagen, 12945). 16 colonies of each sub-library were sequenced for validation (Plasmidsaurus). Sub-libraries corresponding to the same CRE were pooled, and the complete variant library was PCR-amplified and cloned into the MPRA pGL4-GFP backbone vector for downstream assays, as detailed below.

#### MPRA library cloning

20 ng of synthetic oligonucleotide pools corresponding to the CLEM4, C/EBPRE, and IHRR libraries, as well as pooled pUC19 plasmids for the XREM and DRR libraries, were PCR-amplified for 15 cycles using primers that introduced a 5′ overhang compatible with the pGL4-GFP backbone and a 3′ overhang complementary to the CYP3A4 proximal promoter (primers 5-RE-f01 and 5-RE-r01). 20 ng of CYP3A4 proximal promoter was amplified in two successive PCR steps: the first added a spacer sequence at the 3′ end (primers 5-CPP-f01 and 5-CPP-r01), and the second introduced a 15-bp random barcode (primers 5-CPP-f01, 5-CPP-f02). For the promoter library, pooled pUC19 plasmids generated by fragment-based cloning were PCR-amplified to add a 3′ overhang complementary to the CLEM4 enhancer and a 5′ 15-bp barcode (primers 5-CP-lib-f01 and 5-CPP-r01; **Supplementary Data 2**). The CLEM4 reference sequence was PCR-amplified to introduce a 5′ overhang compatible with the pGL4-GFP vector (primers CLEM_F_OH and CLEM_R; **Supplementary Data 2**). The pGL4-GFP backbone was linearized by digestion with *AgeI-HF* and *SacI-HF*. Library inserts, promoter fragments, or CLEM4 enhancer fragment in the case of the promoter library, and linearized vector were purified using 0.8X AMPure XP beads and assembled by Gibson cloning using the NEBuilder HiFi DNA Assembly Cloning Kit (New England Biolabs). The assembly product was again purified with 0.65X Ampure XP beads and transformed into *E. coli NEB 10-beta* electrocompetent cells as described above. The IHRR sequence contains a motif of a prokaryotic promoter, which likely resulted in expression of a toxic peptide in *E. coli* and reduced cloning efficiency. To mitigate this toxicity, the IHRR library was cloned using CopyCutter EPI400 competent cells (LGC Biosearch Technologies, C400EL10), which support low-copy plasmid maintenance.

#### Association sequencing

To establish variant-barcode associations in the MPRA libraries, we performed long-read sequencing of the plasmid pool. CRE-promoter-barcodes were amplified from 2 ug of plasmid in total, in separate reactions for each sub-library (primers pGL4-asso-CRE_F and pGL4-asso-R, sequences in **Supplementary Data 2**) for 7 cycles. The amplicons were purified using 0.6X AMPure XP beads and then digested with *BsrGI or SpeI* for the promoter library (New England Biolabs, R3182) to linearize the contaminant plasmid template. The restriction enzymes were selected to avoid cutting within any of the variants in the library. The amplicons were size-selected using the Blue Pippin system (Sage Science) with the 0.75% Dye-Free Agarose Gel Cassette and External Marker S1 (Sage Science, BLF7510) (Settings: High Pass and low voltage 1-6kb) and 3kb with tight range collection mode to eliminate the contaminant plasmid template. The eluted DNA was purified with 1X AMPure XP beads and quantified using the High-Sensitivity DNA D5000 TapeStation system (Agilent). An amplicon library was generated following standard procedures and sequenced on a PacBio Revio system using one Revio SMRT cell.

#### Library transfection

5ug of each CRE library was transfected into HepG2 cells (1×10^6^ cells in 60mm plates) using X-tremeGENE HP DNA transfection reagent at a reagent:DNA ratio of 2:1, following the manufacturer’s protocol, in three biological replicates. For the XREM library, transfected cells were treated 24 hours post-transfection with 10 µM rifampin. The MPRA sequencing libraries were prepared as described by Gordon^64^. Briefly, three days post infection, cells were washed 3 times with PBS and DNA and RNA were extracted using the Qiagen AllPrep DNA/RNA kit (Qiagen, 80204), according to the manufacturer’s protocol. The RNA barcodes were reverse transcribed using the primer P7-pLSmP-ass16UMI-gfp and Superscript IV Reverse Transcriptase (Invitrogen, 18090010) according to the manufacturer’s protocol. cDNA and DNA barcodes were amplified by PCR for 3 cycles (primers P7-pLSmP-ass16UMI-gfp and P5-pLSmP-5bc-i#) to add sample index and UMI, then purified with 1.4X AMPure XP beads. The barcodes were then amplified using P5 and P7 primers for 10-15 cycles, then purified with 1.2X AMPure XP beads. The MPRA sequencing libraries were sequenced using the Illumina Nextseq High-output using custom primers (Read1: pLSmP-ass-seq-ind1; Read2: pLSmP-bc-seq; Index1: pLSmP-UMI-seq; Index2: pLSmP-5bc-seq-R2) and with the following run settings (Read1, Index1, Index2, Read2): 15, 16, 10, 15.

### Computational analysis

#### Barcode-variant Association

Experiments were sequenced jointly using the PacBio long-read sequencing platform. Reads were demultiplexed based on sequence length into promoter and enhancer experiments. PBMM2 v1.16 (https://github.com/PacificBiosciences/pbmm2), a wrapper for minimap2^65^ was used to generate a mapping index from the libraries’ FASTA files. Following indexing, reads were aligned with PBMM2 using the following options: --preset SUBREAD, --sort, --best-n 1, --min-concordance-perc 100.

For barcode extraction, aligned sequences were scanned for a consistent plasmid sequence adjacent to the barcode “GCAAAGTGAACACATCGCTAAGCGAAAGCTAAG”. The following 15 bases were used as a barcode. Barcodes associated with multiple sequences were dropped from further analysis, and an association file mapping inserts to barcodes was generated for each library.

#### Barcode Count

MPRAsnakeflow v0.5.4^36^ ‘experiment’ step was used to analyze the MPRA RNA and DNA values per barcode per tested sequence, the pipeline was run separately for each library. To quantify variant effects BCalm v0.99^66^ was used to calculate log2 fold change and Benjamini-Hochberg multiple testing adjusted P-values per variant.

#### Differential Analysis

BCalm v0.99 was used to obtain a variant effect quantification (log2 fold change) and a Benjamini-Hochberg adjusted p-value per variant, to identify significant variant effects (adjusted p-value < 0.1). Variants containing less than 10 barcodes were filtered out. Outlier barcodes with an absolute z-score of more than 3 in RNA counts within the same variant were also filtered out.

#### Variant distribution analysis

Spatial uniformity of variants within each CRE was evaluated using a one-sample Kolmogorov-Smirnov test with Benjamini-Hochberg correction across loci. Local enrichment was assessed using 50 bp sliding-window Poisson tests based on each locus’s background variant rate. Windows with BH-adjusted q < 0.05 were considered significant hotspots or coldspots.

#### TFBS motif analysis

JASPAR2026 non-redundant vertebrate position weight matrices (PWMs) were filtered using an expression threshold of TPM>1 in HepG2 for the HepG2 tested libraries, and LS174 for IHRR^67^. To assess variant predicted TFBS effects, variant and reference oligos were scanned with the memelite implementation of FIMO, with a threshold of 0.0001^68^. Motifs were assigned as gains or losses based on FIMO score differences that shifted the score above or below 10, respectively. To annotate motifs in reference CREs, sequences were scanned with FIMO, with a threshold of 0.0005 using a subset of the JASPAR2026 PWMs, containing 34 PWMs of RXR, VDR, RAR, NR1H, NR1I, PPAR, CEBP, HNF1, HNF4, FOXA, and ARNT, and their composite PWMs.

## Acknowledgements

We thank Drs. Sarah Tishkoff, Michael Alan McQuillan and Chao Zhang for providing the data of variants in the WGS of 180 African individuals. We thank Dr. Ofer Yizhar-Barnea for insightful discussions and valuable advice throughout the study. We thank Dr. Ilias Georgakopoulos Soares for assistance with retrieving and curating the cancer-associated data. This work was supported in part by the National Human Genome Research Institute (NHGRI) grant numbers UM1HG009408 (N.A.) and UM1HG011966 (N.A.) and the UCSF Mary Anne Koda-Kimble Seed Award for Innovation (N.A.).

## Author contributions

Y.G. and N.A. conceived the study and devised the methodology; Y.G. carried out experimental work; B.K. performed data analysis; N.A. supervised the project and acquired funding. Y.G, B.K. and N.A. wrote, reviewed and edited the paper with contributions from all authors.

## Data availability

MPRA datasets have been deposited to the Impact of Genomic Variation on Function consortium (IGVF) database under the following accessions, IGVFDS8965KQTP for C/EBPRE, IGVFDS1949NXAF for CLEM4, IGVFDS6913MGLB for DRR, IGVFDS5092STSN for IHRR, IGVFDS8542QKRX for Promoter, and IGVFDS0532KPPJ for XREM.

Detailed protocol is available at https://www.protocols.io/blind/565F66AD1B5511F18EC00A58A9FEAC02

## Code availability

Code for the long-read MPRA association stage is available at https://github.com/Ahituv-lab/CypMPRA and on Zenodo at https://zenodo.org/records/19363634.

## References

1. Johnson, G. D. et al. Human genome-wide measurement of drug-responsive regulatory activity. Nat Commun 9, 5317 (2018).

2. Chatterjee, S. & Ahituv, N. Gene Regulatory Elements, Major Drivers of Human Disease. Annu Rev Genomics Hum Genet 18, 45–63 (2017).

3. Luizon, M. R. & Ahituv, N. Uncovering Drug-Responsive Regulatory Elements. Pharmacogenomics 16, 1829–1841 (2015).

4. Ward, L. D. & Kellis, M. Interpreting noncoding genetic variation in complex traits and human disease. Nat Biotechnol 30, 1095–1106 (2012).

5. Lewis, D. F. V. & Ito, Y. Human CYPs involved in drug metabolism: structures, substrates and binding affinities. Expert Opin Drug Metab Toxicol 6, 661–674 (2010).

6. Paine, M. F. et al. The human intestinal cytochrome P450 ‘pie’. Drug Metab Dispos 34, 880–886 (2006).

7. Zanger, U. M. & Schwab, M. Cytochrome P450 enzymes in drug metabolism: Regulation of gene expression, enzyme activities, and impact of genetic variation. Pharmacology & Therapeutics 138, 103–141 (2013).

8. Dobon, B., Rossell, C., Walsh, S. & Bertranpetit, J. Is there adaptation in the human genome for taste perception and phase I biotransformation? BMC Evolutionary Biology 19, 39 (2019).

9. McArthur, A. G. et al. Phylogenetic Analysis of the Cytochrome P450 3 (CYP3) Gene Family. J Mol Evol 57, 200–211 (2003).

10. Richard-St-Hilaire, A. et al. Signatures of Co-evolution and Co-regulation in the CYP3A and CYP4F Genes in Humans. Genome Biol Evol 16, evad236 (2024).

11. Qiu, H. et al. CYP3 phylogenomics: evidence for positive selection of CYP3A4 and CYP3A7. Pharmacogenetics and Genomics 18, 53 (2008).

12. Kasarla, S. S., Garikapati, V., Kumar, Y. & Dodoala, S. Interplay of Vitamin D and CYP3A4 Polymorphisms in Endocrine Disorders and Cancer. Endocrinol Metab 37, 392–407 (2022).

13. Stipp, M. C. & Acco, A. Involvement of cytochrome P450 enzymes in inflammation and cancer: a review. Cancer Chemother Pharmacol 87, 295–309 (2021).

14. Attia, H. R. M. et al. CYP2C8 rs11572080 and CYP3A4 rs2740574 risk genotypes in paclitaxel-treated premenopausal breast cancer patients. Sci Rep 14, 7922 (2024).

15. Martínez-Jiménez, C. P., Gómez-Lechón, M. J., Castell, J. V. & Jover, R. Transcriptional Regulation of the Human Hepatic CYP3A4: Identification of a New Distal Enhancer Region Responsive to CCAAT/Enhancer-Binding Protein β Isoforms (Liver Activating Protein and Liver Inhibitory Protein). Mol Pharmacol 67, 2088–2101 (2005).

16. Bertilsson, G., Berkenstam, A. & Blomquist, P. Functionally Conserved Xenobiotic Responsive Enhancer in Cytochrome P450 3A7. Biochem. Biophys. Res. Commun. 280, 139–144 (2001).

17. Goodwin, B., Hodgson, E. & Liddle, C. The orphan human pregnane X receptor mediates the transcriptional activation of CYP3A4 by rifampicin through a distal enhancer module. Molecular pharmacology 56, 1329–1339 (1999).

18. Matsumura, K. et al. Identification of a Novel Polymorphic Enhancer of the Human CYP3A4 Gene. Mol Pharmacol 65, 326–334 (2004).

19. Collins, J. M. et al. Regulatory variants in a novel distal enhancer regulate the expression of CYP3A4 and CYP3A5. Clinical and Translational Science 15, 2720–2731 (2022).

20. Tegude, H. et al. Molecular Mechanism of Basal CYP3A4 Regulation by Hepatocyte Nuclear Factor 4α: Evidence for Direct Regulation in the Intestine. Drug Metab Dispos 35, 946–954 (2007).

21. Inoue, F. & Ahituv, N. Decoding enhancers using massively parallel reporter assays. Genomics 106, 159–164 (2015).

22. Kircher, M. et al. Saturation mutagenesis of twenty disease-associated regulatory elements at single base-pair resolution. Nat Commun 10, 3583 (2019).

23. Patwardhan, R. P. et al. High-resolution analysis of DNA regulatory elements by synthetic saturation mutagenesis. Nat Biotechnol 27, 1173–1175 (2009).

24. Melnikov, A. et al. Systematic dissection and optimization of inducible enhancers in human cells using a massively parallel reporter assay. Nat Biotechnol 30, 271–277 (2012).

25. Martinez-Ara, M., Comoglio, F., van Arensbergen, J. & van Steensel, B. Systematic analysis of intrinsic enhancer-promoter compatibility in the mouse genome. Mol Cell 82, 2519–2531.e6 (2022).

26. Bergman, D. T. et al. Compatibility rules of human enhancer and promoter sequences. Nature 607, 176–184 (2022).

27. Nebert, D. W. & Russell, D. W. Clinical importance of the cytochromes P450. Lancet 360, 1155–1162 (2002).

28. Karczewski, K. J. et al. The mutational constraint spectrum quantified from variation in 141,456 humans. Nature 581, 434–443 (2020).

29. The Cancer Genome Atlas Program (TCGA) - NCI. https://www.cancer.gov/ccg/research/genome-sequencing/tcga (2022).

30. Reich, D. et al. Genetic history of an archaic hominin group from Denisova Cave in Siberia. Nature 468, 1053–1060 (2010).

31. Prüfer, K. et al. The complete genome sequence of a Neanderthal from the Altai Mountains. Nature 505, 43–49 (2014).

32. Arzumanian, V. A., Kiseleva, O. I. & Poverennaya, E. V. The Curious Case of the HepG2 Cell Line: 40 Years of Expertise. International Journal of Molecular Sciences 22, 13135 (2021).

33. Chiang, T.-S., Yang, K.-C., Chiou, L.-L., Huang, G.-T. & Lee, H.-S. Enhancement of CYP3A4 Activity in Hep G2 Cells by Lentiviral Transfection of Hepatocyte Nuclear Factor-1 Alpha. PLOS ONE 9, e94885 (2014).

34. Sebastian, T. & Johnson, P. F. Stop and Go: Anti-Proliferative and Mitogenic Functions of the Transcription Factor C/EBPβ. Cell Cycle 5, 953–957 (2006).

35. Klein, J. C. et al. A systematic evaluation of the design and context dependencies of massively parallel reporter assays. Nat Methods 17, 1083–1091 (2020).

36. Rosen, J. D. et al. Uniform processing and analysis of IGVF massively parallel reporter assay data with MPRAsnakeflow. 2025.09.25.678548 Preprint at 10.1101/2025.09.25.678548 (2025).

37. Chen, S. et al. A genome-wide mutational constraint map quantified from variation in 76,156 human genomes. 2022.03.20.485034 Preprint at 10.1101/2022.03.20.485034 (2022).

38. Hudson (Chairperson), T. J., et al. International network of cancer genome projects. Nature 464, 993–998 (2010).

39. Sadeghi, H., Hashemnia, V., Nazemalhosseini-Mojarad, E., Ghasemi, M. R. & Mirfakhraie, R. Correlated downregulation of VDR and CYP3A4 in colorectal cancer. Mol Biol Rep 50, 1385–1391 (2023).

40. Wang, F. et al. Activation/Inactivation of Anticancer Drugs by CYP3A4: Influencing Factors for Personalized Cancer Therapy. Drug Metabolism and Disposition 51, 543–559 (2023).

41. Sumantran, V. N., Mishra, P., Bera, R. & Sudhakar, N. Microarray Analysis of Differentially-Expressed Genes Encoding CYP450 and Phase II Drug Metabolizing Enzymes in Psoriasis and Melanoma. Pharmaceutics 8, 4 (2016).

42. Biggs, J. S. et al. Transcription Factor Binding to a Putative Double E-Box Motif Represses CYP3A4 Expression in Human Lung Cells. Molecular Pharmacology 72, 514–525 (2007).

43. Ballas, N., Grunseich, C., Lu, D. D., Speh, J. C. & Mandel, G. REST and its corepressors mediate plasticity of neuronal gene chromatin throughout neurogenesis. Cell 121, 645–657 (2005).

44. Sohn, M. et al. Whole exome sequencing for the identification of CYP3A7 variants associated with tacrolimus concentrations in kidney transplant patients. Sci Rep 8, 18064 (2018).

45. Le lay, J. & Kaestner, K. H. The Fox Genes in the Liver: From Organogenesis to Functional Integration. Physiological Reviews 90, 1–22 (2010).

46. Huggins, C. J. et al. C/EBPγ Is a Critical Regulator of Cellular Stress Response Networks through Heterodimerization with ATF4. Molecular and Cellular Biology 36, 693–713 (2016).

47. Oliva-Vilarnau, N. et al. Comparative analysis of YAP/TEAD inhibitors in 2D and 3D cultures of primary human hepatocytes reveals a novel non-canonical mechanism of CYP induction. Biochem Pharmacol 215, 115755 (2023).

48. Azuma, K., Urano, T., Watabe, T., Ouchi, Y. & Inoue, S. PROX1 suppresses vitamin K-induced transcriptional activity of steroid and xenobiotic receptor. Genes to Cells 16, 1063–1070 (2011).

49. Jin, H. et al. Systematic transcriptional analysis of human cell lines for gene expression landscape and tumor representation. Nat Commun 14, 5417 (2023).

50. Zheng, Y. et al. CYP3A4*1B Polymorphism and Cancer Risk: A Meta-Analysis Based on 55 Case-control Studies. Ann Clin Lab Sci 48, 538–545 (2018).

51. Shi, W.-L., Tang, H.-L. & Zhai, S.-D. Effects of the CYP3A4*1B Genetic Polymorphism on the Pharmacokinetics of Tacrolimus in Adult Renal Transplant Recipients: A Meta-Analysis. PLOS ONE 10, e0127995 (2015).

52. Ettienne, E. B. et al. Pharmacogenomics-guided policy in opioid use disorder (OUD) management: An ethnically-diverse case-based approach. Addictive Behaviors Reports 6, 8–14 (2017).

53. Lamba, J. K., Lin, Y. S., Schuetz, E. G. & Thummel, K. E. Genetic contribution to variable human CYP3A-mediated metabolism. Advanced Drug Delivery Reviews 54, 1271–1294 (2002).

54. Spurdle, A. B. et al. TheCYP3A4*1Bpolymorphism has no functional significance and is not associated with risk of breast or ovarian cancer. Pharmacogenetics and Genomics 12, 355 (2002).

55. Gaedigk, A., Casey, S. T., Whirl-Carrillo, M., Miller, N. A. & Klein, T. E. Pharmacogene Variation Consortium: A Global Resource and Repository for Pharmacogene Variation. Clin Pharmacol Ther 110, 542–545 (2021).

56. Klein, K. & Zanger, U. M. Pharmacogenomics of Cytochrome P450 3A4: Recent Progress Toward the “Missing Heritability” Problem. Front. Genet. 4, (2013).

57. Wojnowski, L. & Kamdem, L. K. Clinical implications of CYP3A polymorphisms. Expert Opinion on Drug Metabolism & Toxicology 2, 171–182 (2006).

58. Werk, A. N. & Cascorbi, I. Functional Gene Variants of CYP3A4. Clinical Pharmacology & Therapeutics 96, 340–348 (2014).

59. Zanger, U. M. et al. Genetics, Epigenetics, and Regulation of Drug-Metabolizing Cytochrome P450 Enzymes. Clin. Pharmacol. Ther. 95, 258–261 (2014).

60. Kumarakulasingham, M. et al. Cytochrome P450 Profile of Colorectal Cancer: Identification of Markers of Prognosis. Clin Cancer Res 11, 3758–3765 (2005).

61. Rohs, R. et al. The role of DNA shape in protein–DNA recognition. Nature 461, 1248–1253 (2009).

62. Hinrichs, A. S. et al. The UCSC Genome Browser Database: update 2006. Nucleic Acids Res 34, D590–D598 (2006).

63. Fan, S. et al. Whole-genome sequencing reveals a complex African population demographic history and signatures of local adaptation. Cell 186, 923–939.e14 (2023).

64. lentiMPRA and MPRAflow for high-throughput functional characterization of gene regulatory elements | Nature Protocols. https://www.nature.com/articles/s41596-020-0333-5.

65. Minimap2: pairwise alignment for nucleotide sequences | Bioinformatics | Oxford Academic. https://academic.oup.com/bioinformatics/article/34/18/3094/4994778.

66. Keukeleire, P. et al. Using individual barcodes to increase quantification power of massively parallel reporter assays. BMC Bioinformatics 26, 52 (2025).

67. Klijn, C. et al. A comprehensive transcriptional portrait of human cancer cell lines. Nat Biotechnol 33, 306–312 (2015).

68. Grant, C. E., Bailey, T. L. & Noble, W. S. FIMO: scanning for occurrences of a given motif. Bioinformatics 27, 1017–1018 (2011).

