## Supplemental Figure 1-5 for "Massively parallel reporter assays for *CYP3A4* enhancer variants alongside their native promoter"

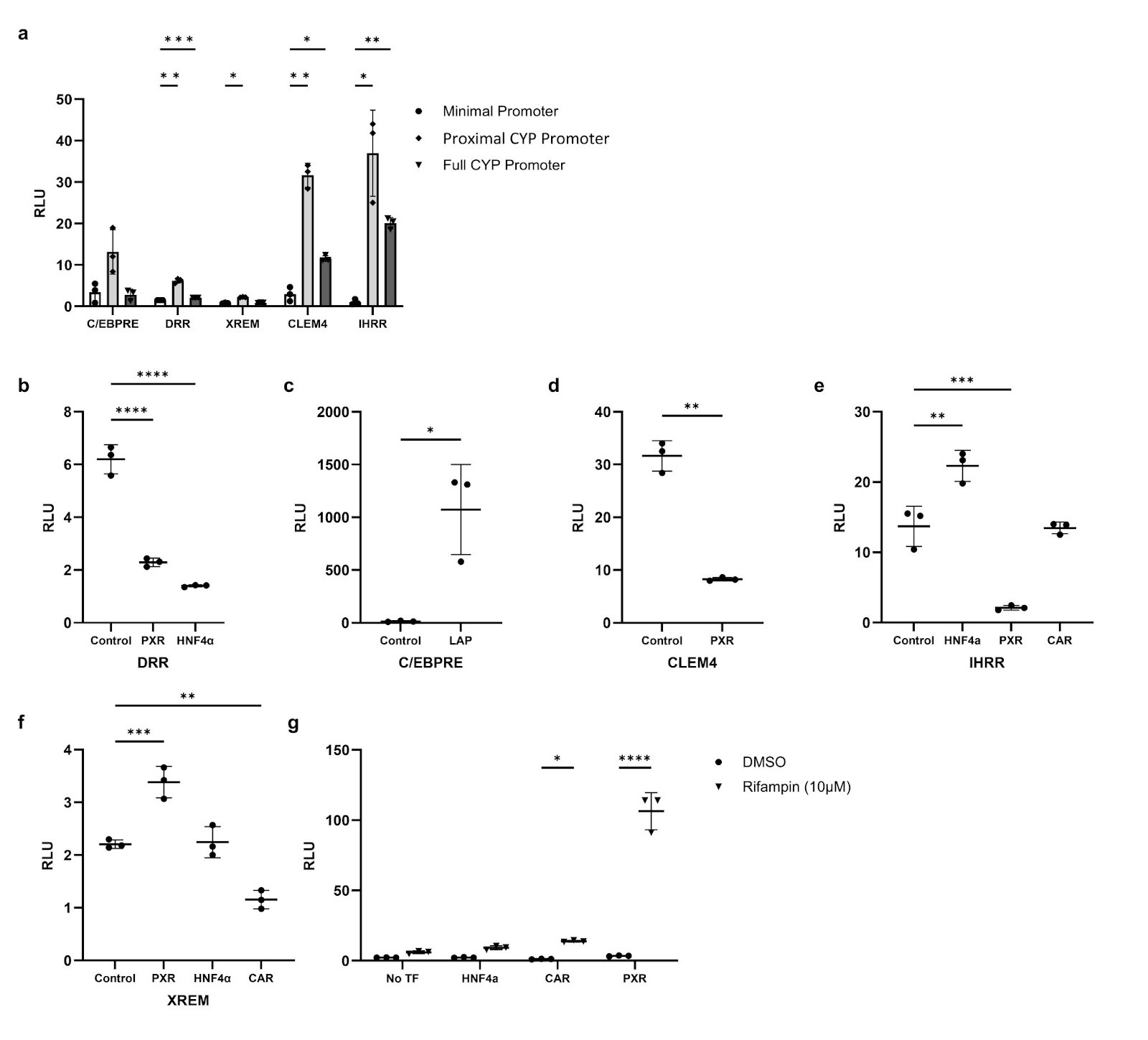

**Supplementary Fig. 1: Luciferase reporter assays to determine promoter context and transcription factor (TF) requirements of *CYP3A4* regulatory elements.** (**a**) Activity of reference *CYP3A4* enhancers cloned upstream of a minimal promoter, the proximal *CYP3A4* promoter, or the full-length *CYP3A4* promoter in HepG2 cells. (**b–f**) Enhancer activity following co-expression of TFs known to regulate *CYP3A4* expression. (**g**) XREM induction upon rifampin treatment in the presence of TFs. Data are shown as individual replicates with mean ± s.d. RLU, relative light units. Statistical significance is indicated (*P < 0.05; **P < 0.01; ***P < 0.001; ****P < 0.0001).

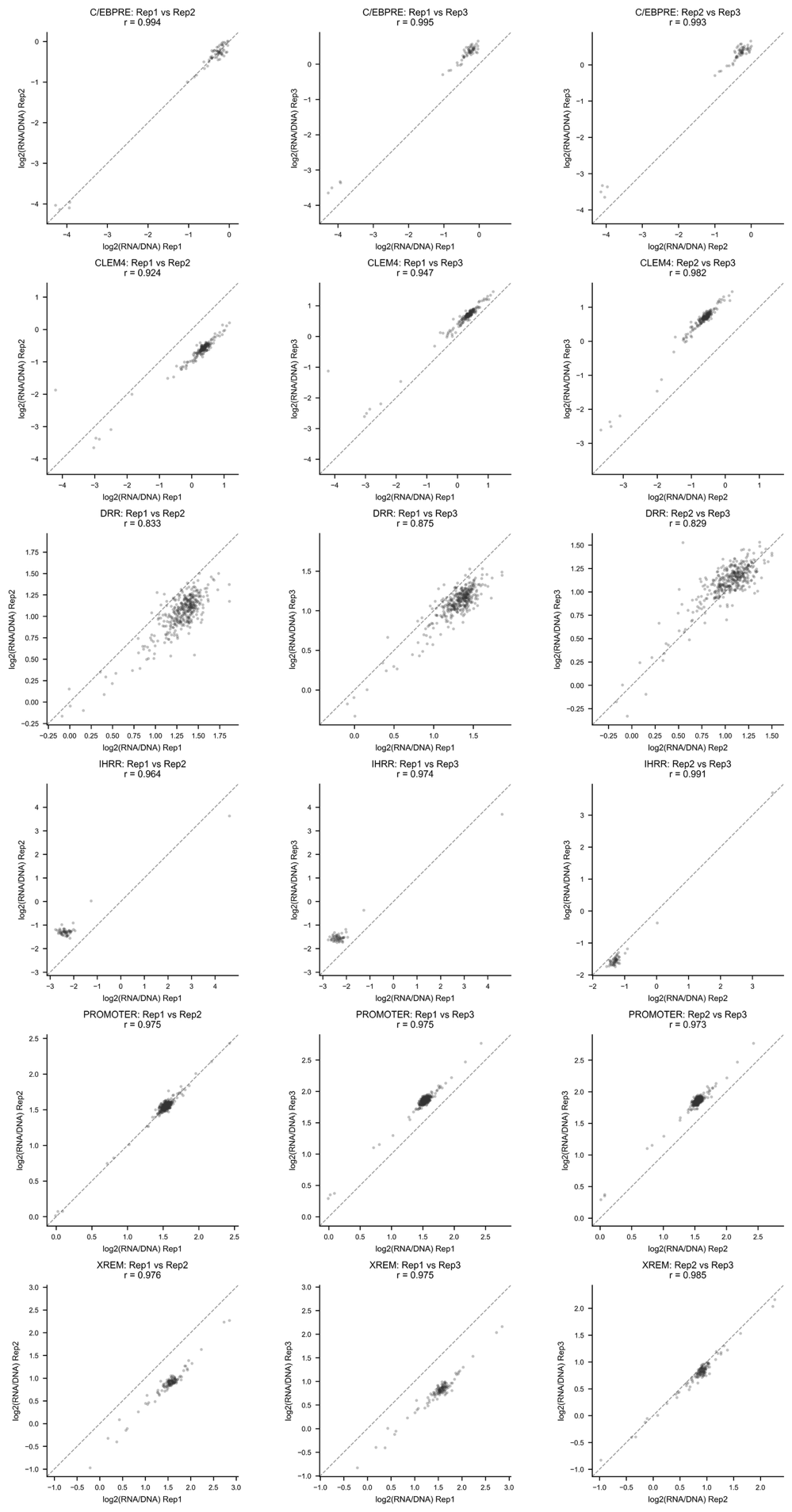

**Supplementary Fig. 2: Replicate correlation.** Scatter plots show Pearson correlations between replicates for the average log2(RNA/DNA) of barcodes for assayed oligonucleotides.

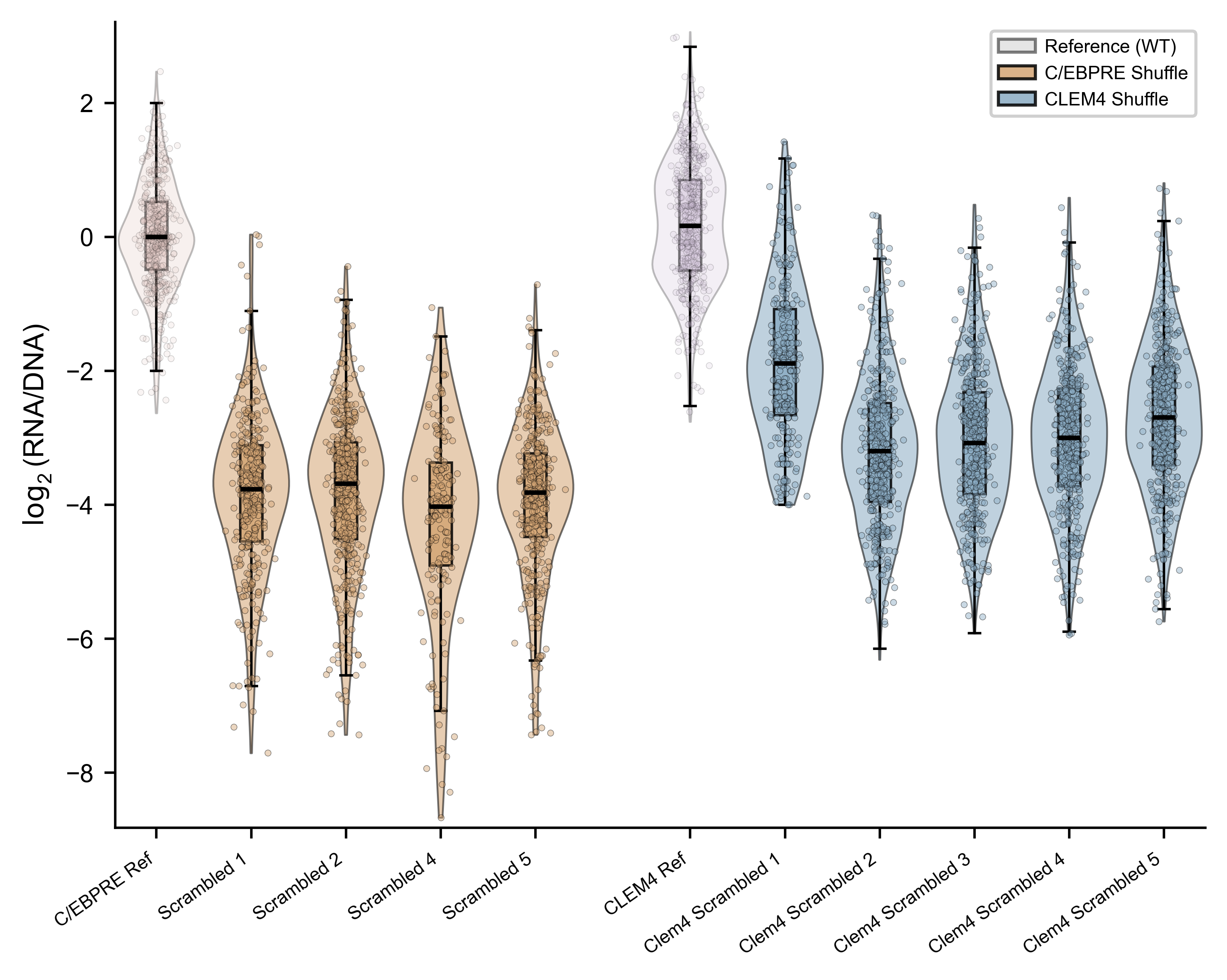
**Supplementary Fig. 3: Scramble controls:** Violin plots show per-barcode log₂(RNA/DNA) activity for reference and dinucleotide shuffled sequence controls.

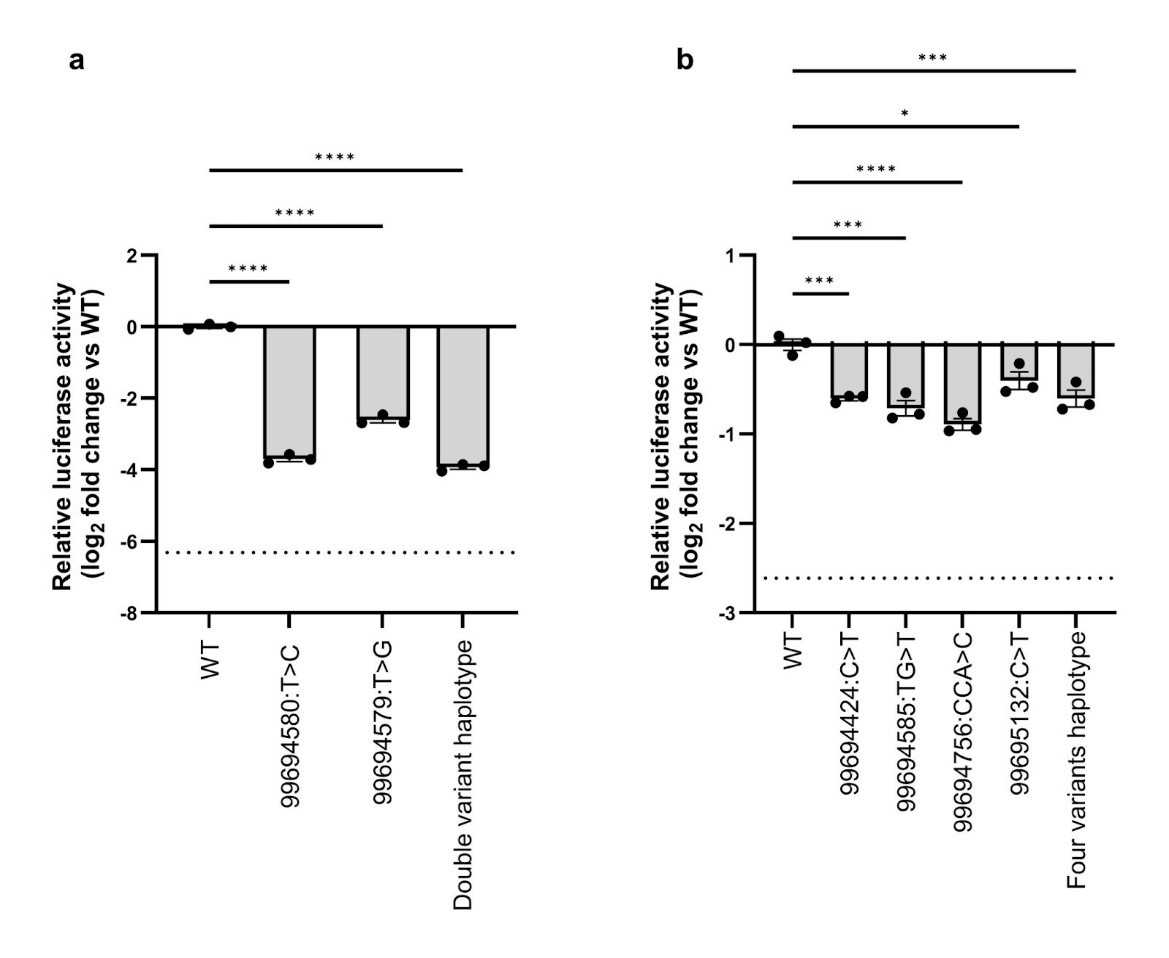

**Supplementary Fig. 4: Luciferase activity of individual variants and the combined haplotypes.**

Data were log_2_-transformed fold change relative to wild type. The dashed line indicates the expected additive effect of single variants. Points represent biological replicates; bars denote mean ± SEM. (**a**) P-values were calculated by two-way ANOVA with an interaction term. P(interaction)<0.0001. (**b**) P-values were calculated by a two-tailed unpaired t-test comparing observed versus expected additive activity. *P < 0.05, **P < 0.01, ***P < 0.001, ****P < 0.0001.

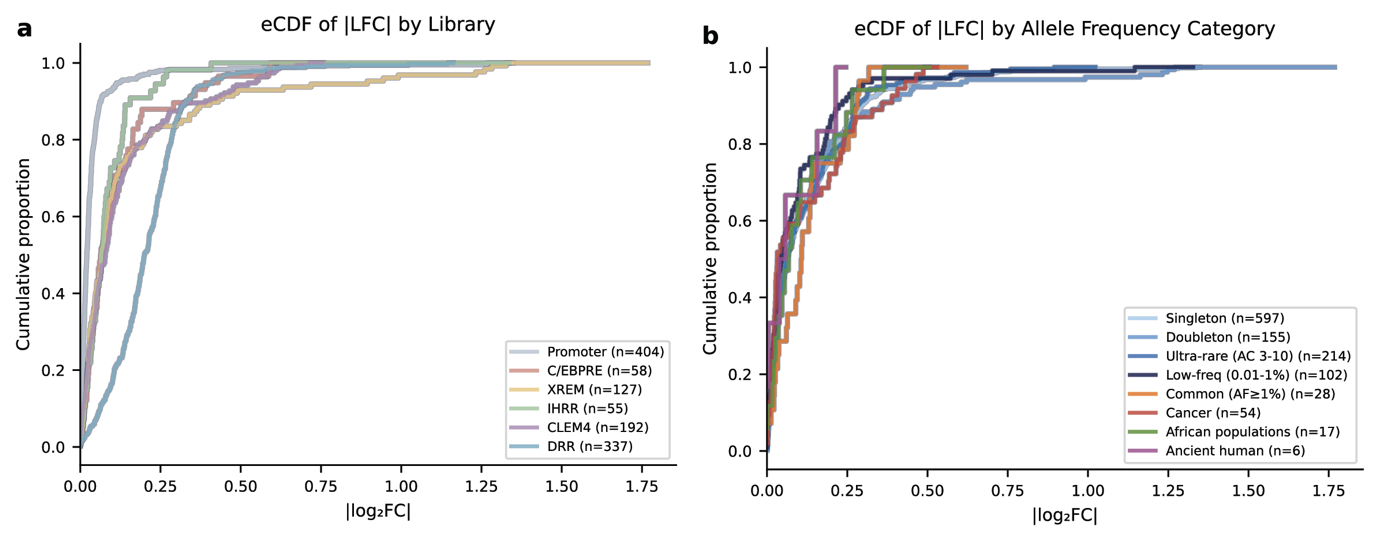

**Supplementary Fig. 5: Effects of *CYP3A4* CRE variants in the MPRA.** Cumulative distribution functions showing the frequencies of variants with specific MPRA log_2_FoldChange measurements relative to the reference, across different categories. (**a**) Stratified by CRE. (**b**) Stratified by variant characterization.

**Supplementary Data 1**: All assayed variants, with allele frequencies, BCalm, and FIMO scores

**Supplementary Data 2:** List of primers

**Supplementary Data 3:** Plasmid maps
